# Antiviral face mask functionalized with solidified hand soap: low-cost infection prevention clothing against enveloped viruses such as SARS-CoV-2

**DOI:** 10.1101/2021.08.13.456326

**Authors:** Alba Cano-Vicent, Alberto Tuñón-Molina, Miguel Martí, Yukiko Muramoto, Takeshi Noda, Kazuo Takayama, Ángel Serrano-Aroca

**Author notes:** These authors contributed equally to this work (Alba Cano-Vicent, Alberto Tuñón-Molina). **Corresponding authors**: **Kazuo Takayama** - Center for iPS Cell Research and Application, Kyoto University, Kyoto 606-8397, Japan; (K.T); **Ángel Serrano-Aroca** - Biomaterials and Bioengineering Lab, Centro de Investigación Traslacional San Alberto Magno, Universidad Católica de Valencia San Vicente Mártir, c/Guillem de Castro 94, Valencia 46001, Spain; (Á.S-A). **Alba Cano-Vicent** -, **Alberto Tuñón-Molina** -, **Miguel Martí** -, **Yukiko Muramoto** -, **Takeshi Noda**.

## Abstract

Infection prevention clothing is becoming an essential protective tool in the current pandemic, especially because now we know that SARS-CoV-2 can easily infect humans in poorly ventilated indoor spaces. However, commercial infection prevention clothing is made of fabrics that are not capable of inactivating the virus. Therefore, viral infections of symptomatic and asymptomatic individuals wearing protective clothing such as masks can occur through aerosol transmission or by contact with the contaminated surfaces of the masks, which are suspected as an increasing source of highly infectious biological waste. Herein, we report an easy fabrication method of a novel antiviral non-woven fabric containing polymer filaments that were coated with solidified hand soap. This extra protective fabric is capable of inactivating enveloped viruses such as SARS-CoV-2 and phi 6 in one minute of contact. In this study, this antiviral fabric was used to fabricate an antiviral face mask and did not show any cytotoxic effect in human keratinocyte HaCaT cells. Furthermore, this antiviral non-woven fabric could be used for the fabrication of other infection prevention clothing such as caps, scrubs, shirts, trousers, disposable gowns, overalls, hoods, aprons, and shoe covers. Therefore, this low-cost technology could provide a wide range of infection protective tools to combat COVID-19 and future pandemics in developed and underdeveloped countries.

## 1. INTRODUCTION

The current coronavirus disease 2019 (COVID-19) pandemic has globally spread around more than 200 countries since December 2019.^1–5^ This new disease is caused by the Severe Acute Respiratory Syndrome Coronavirus 2 (SARS-CoV-2), which is an enveloped positive-sense single-stranded RNA virus^6,7^ classified in the IV Baltimore group.^8^ SARS-CoV-2 continues to globally spread in spite of the strict lockdowns conducted in many countries and the hot season that did not succeed to flatten the global epidemic curve.^9,10^ SARS-CoV-2 has shown stability from some hours to days in different environmental ambients such as aerosols and diverse types of surfaces: metallic, plastic or cardboard.^11–15^ Therefore, SARS-CoV-2 infections can mainly occur in humans by two main routes: contact with contaminated surfaces or by aerosol transmission, especially in poorly ventilated indoor places.^11–15^ In fact, many studies have shown clear evidence that indoor aerosol transmission is one of the most important transmission ways of SARS-CoV-2, especially through asymptomatic carriers.^16–18^ Thus, many countries have made mandatory wearing respiratory face masks covering mouth and nose as a demonstrated strategy to fight against SARS-CoV-2.^19–22^ However, commercial face mask fabrics can only partially block the viral particles and are not able to inactivate them when they reach the fabric. Thus, the new generation face mask fabrics must be developed with antimicrobial materials^23–30^ capable of inactivating viruses to increase human protection even more. In this regard, several antiviral face mask materials against SARS-CoV-2 have been recently proposed. Although, all of these studies propose expensive materials such as graphene,^31,32^ graphene oxide and polydopamine,^33^ titanium oxide,^34^ zinc,^35^ copper, ^36^ silver,^37^ conjugated polymers and oligomers,^38^ and advanced surface coatings^39–42^ that are designed to inactivate the virus. These antiviral composites are produced with complex and costly manufacturing processes, which render them non-viable for a global solution of the current COVID-19 pandemic strongly affecting both developed and underdeveloped countries. Alternative virucidal compounds such as ethanol (70%), povidone-iodine (7.5%), chloroxylenol (0.05%), chlorhexidine (0.05%), benzalkonium chloride (0.1%) and hand soap solution (1:49) have shown antiviral activity against SARS-CoV-2.^15^ From all these virucidal compounds, the Centers for Disease Control and Prevention has repeatedly recommended that the best and most economical way to prevent the COVID-19 spread of infections and decrease the risk of getting sick is by washing often with hand soap and water.^43^ Furthermore, several commonly available healthcare products have shown virucidal activity with respect to human coronavirus 229e (HCoV-229e), used as a surrogate of SARS-CoV-2, such as Johnson’s Baby shampoo from Johnson & Johnson Consumer Inc and Crest Pro-Health toothpaste from Procter & Gamble.^44^ These healthcare commercial products are composed of many active and inactive ingredients. In this regard, we hypothesized here that a very low amount of commercial hand soap could be physically adsorbed and solidified onto the surfaces of the polymer filaments of a non-woven commercial fabric by the dip-coating method.^45^ Thus, we expect that the absorbed hand soap could form a biofunctional coating capable of inactivating SARS-CoV-2. The study also aims to use this antiviral fabric to fabricate an antiviral face mask with non-cytotoxic effects for human keratinocyte cells. Non-woven fabrics are currently manufactured very rapidly and at a very low price, are flexible and lightweight but resilient, provide good breathability and bacterial filtration, and are hygienic as they are made for single-use.^46^ Therefore, we expect to provide here a global solution by developing an antiviral face mask non-woven fabric using low-cost materials that could contribute to reduce the rate of COVID-19 infections. Furthermore, we expect that this antiviral non-woven fabric could be used for the fabrication of other infection prevention clothing such as caps, scrubs, shirts, trousers, disposable gowns, overalls, hoods, aprons, and shoe covers.

## 2. RESULTS AND DISCUSSION

### 2.1. Fabric morphology and characterization

The electron microscopy morphology of the porous commercial non-woven is shown in Figure 1.

**Figure 1.**
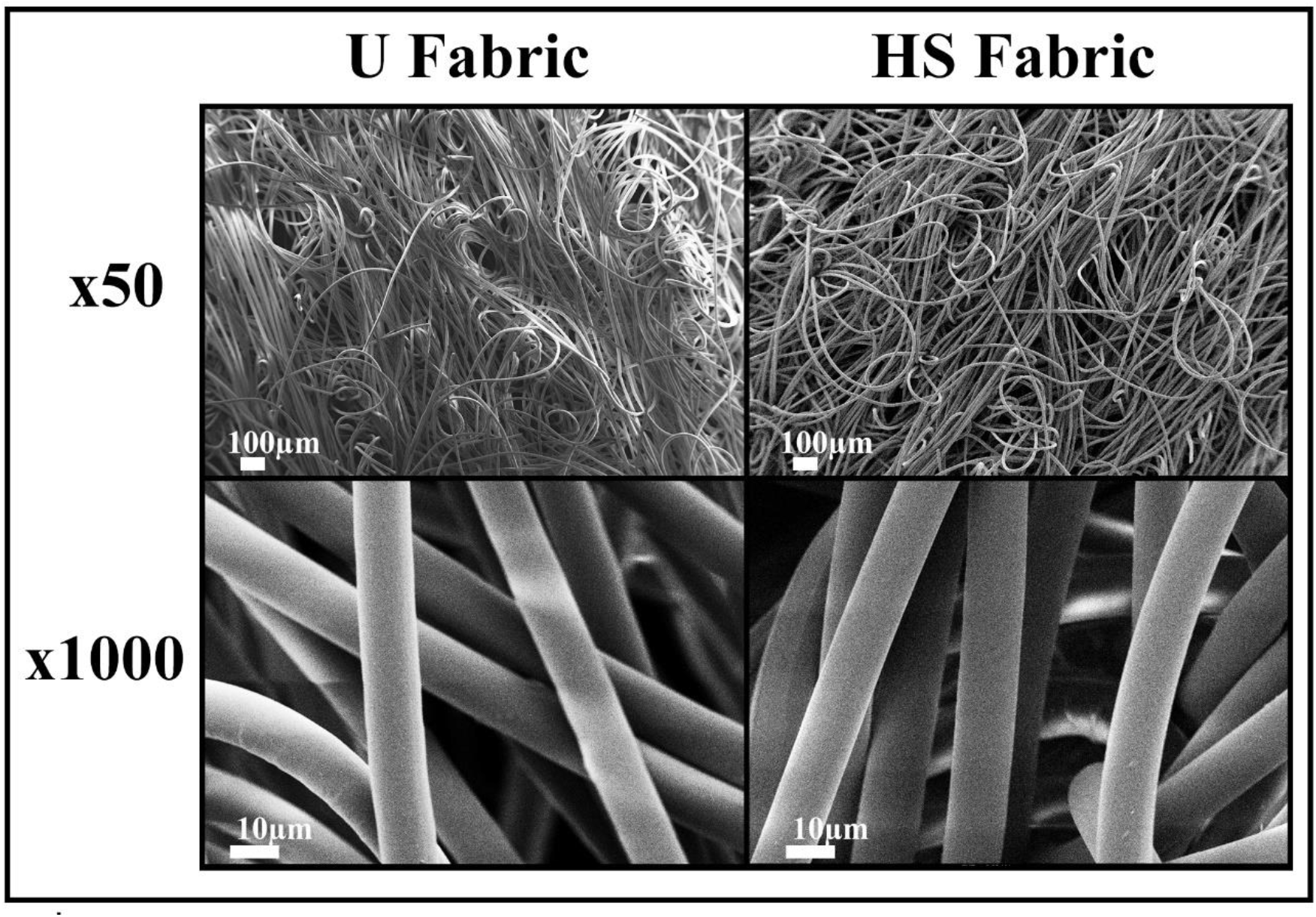
Field emission electron micrographs of the non-woven fabrics. Images of the untreated fabric (U Fabric) and fabric with 0.57±0.03% *w/w* of a biofunctional solidified commercial hand soap (HS Fabric) at two magnifications (×50 and ×1000).

The electron micrographs (Figure 1) and macroscopic images (Figure 2(a)) of the porous non-woven HS fabric do not show any sign of morphological change after the dip coating process performed on the U Fabric.

**Figure 2.**
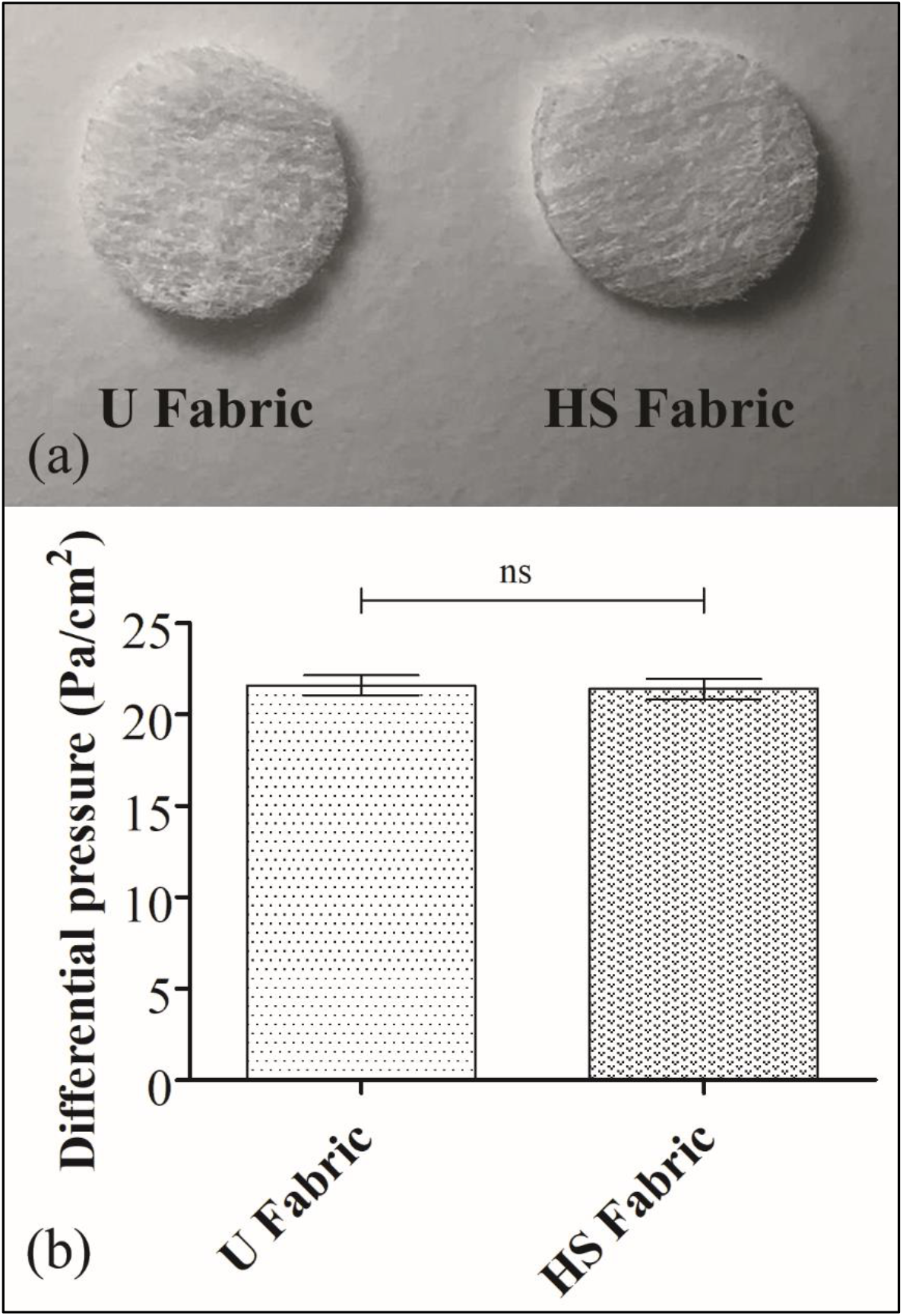
Photographs and differential pressure of the non-woven fabrics: (a) photographs and differential pressure (b) of the untreated fabric (U Fabric) and fabric with 0.57±0.03% *w/w* of a biofunctional solidified commercial hand soap (HS Fabric); ns, not significant.

Therefore, no change of breathability nor bacterial filtration efficiency are expected. In fact, no statistically significant difference of differential pressure (breathability) was found between the U Fabric and the HS Fabric (Figure 2(b)). Furthermore, the developed HS Fabric showed a differential pressure value that falls in the acceptance level (<40 Pa/cm^2^) to be used in the fabrication of face masks according to the standard EN14683:2019+AC:2019.

### 2.2. Antiviral tests

#### 2.2.1. Tests with the biosafe viral model of SARS-CoV-2

We have recently validated the use of bacteriophage phi 6 as a biosafe viral model of SARS-CoV-2 against benzalkonium chloride.^47^ Phage phi 6 is an enveloped double-stranded RNA bacteriophage that belongs to the group III of the Baltimore classification^8^ and it possesses three-part, segmented genome with a total length of ~13.5 kb. Thus, this bacteriophage is also proposed here as biosafe virus model of SARS-CoV-2 against hand soap. RNA extraction and quantification of the phi 6 virus after being in contact with the untreated (U Fabric) and treated (HS Fabric) materials were performed to demonstrate that the virus does not remain trapped in the pores before the antiviral assays, which could give false results. Thus, Figure 3 shows how the amount of RNA did not significantly decrease after being in contact with the HS Fabric or U Fabric with respect to control (same amount of virus without being in contact with the fabrics).

**Figure 3.**
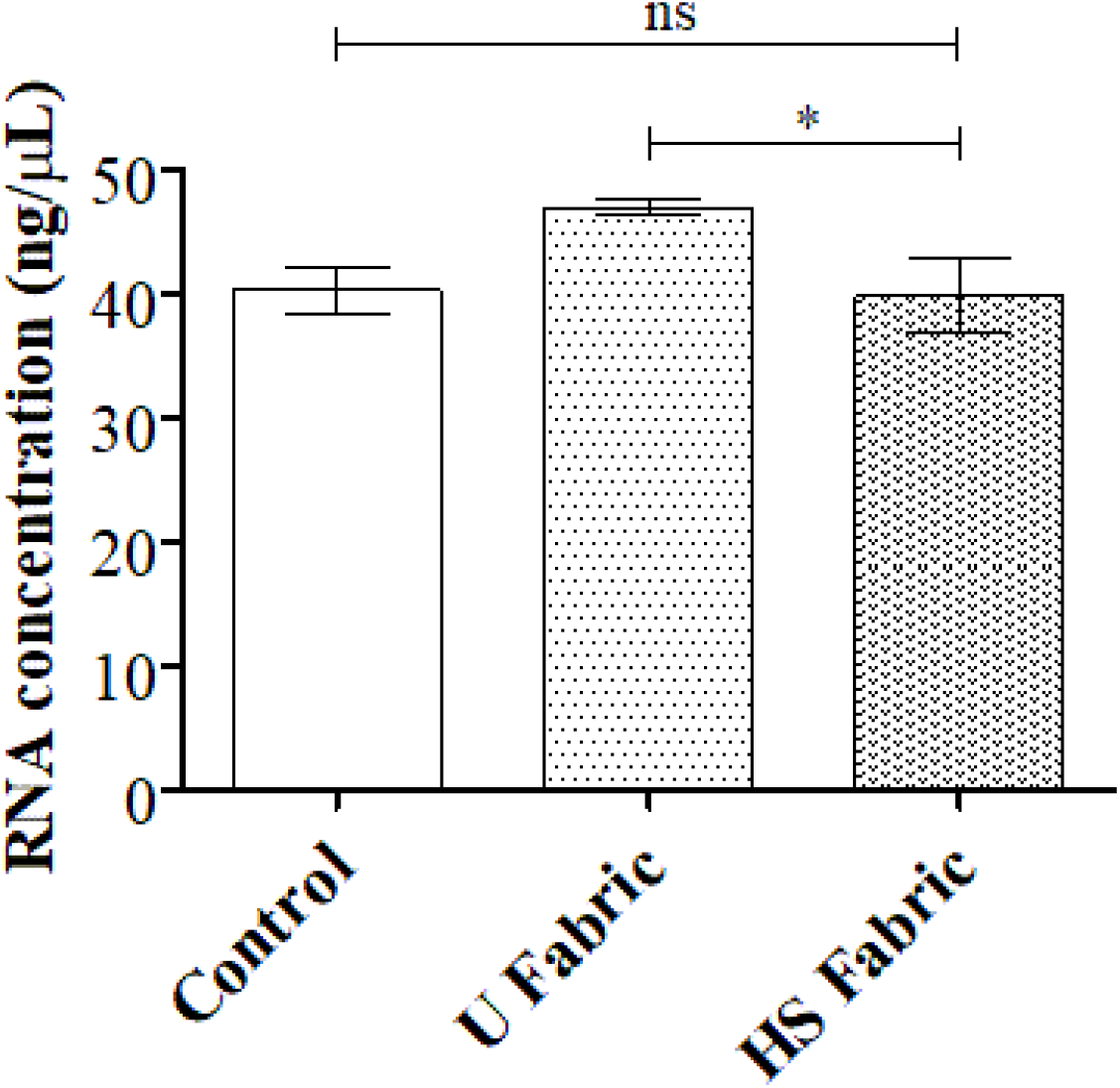
RNA extraction and quantification of the phi 6 virus: RNA concentration in ng/μL of the phi 6 virus measured in the control (without being in contact with the fabrics) and the same amount of virus after being in contact with the U Fabric and HS Fabric for 5 minutes; * *p* > 0.05; ns, not significant.

Therefore, all the viral particles are released from the fabrics after sonication and vortexing. The results of the antiviral tests showed that the HS fabric possesses potent antiviral activity (100% of viral inhibition, see Figure 4) even with just 1 minute of contact with the bacteriophage phi6.

**Figure 4.**
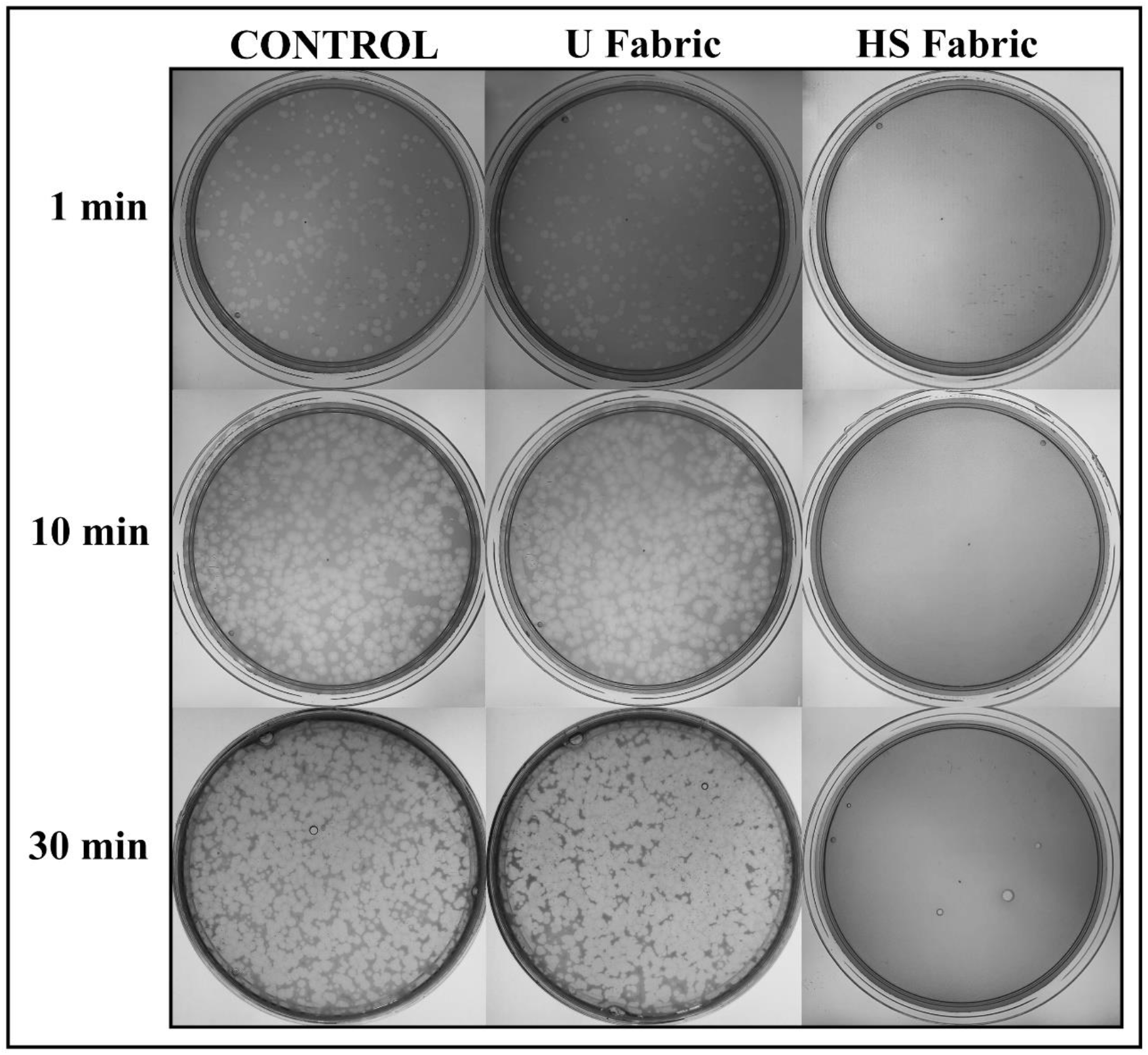
Loss of phage phi 6 viability measured by the double-layer method: undiluted results without being in contact with any fabric (control) and after being in contact with the untreated polymer fabrics (U Fabric), and with the fabric functionalized with solidified commercial hand soap (HS Fabric) during 1, 10 and 30 minutes.

Thus, after 1, 10 or 30 minutes of contact between the HS fabric and the SARS-CoV-2 viral model, bacterial lawns grew in the plate without plaques (see Figure 4). In contrast, the U Fabric showed similar results to control (without being in contact with any fabric) of no antiviral activity as expected. The logarithm of plaque-forming units per mL (PFU/mL) of bacteriophage phi6 after being in contact with the HS Fabric is shown and compared with the U Fabric and control in Table 1.

**Table 1.** Infection titers obtained by the double layer method for the antiviral assay performed with phi 6 and by the TCID50/mL method for SARS-CoV-2 expressed as mean±standard deviation, and percentage of viral inactivation with respect to control without being in contact with any fabric (control), and after being in contact with the untreated fabric (U Fabric), and the fabric with the solidified hand soap coating (HS Fabric) for 1, 10 and 30 minutes.

| Sample | Phi 6 at 1 min (PFU/mL) | Inactivation at 1 min | Phi 6 at 10 min (PFU/mL) | Inactivation at 10 min | Phi 6 at 30 min (PFU/mL) | Inactivation at 30 min |
| --- | --- | --- | --- | --- | --- | --- |
| Control | $1.6 \times 10^5 \pm 1.4 \times 10^4$ | - | $3.2 \times 10^6 \pm 8.9 \times 10^5$ | - | $1.4 \times 10^7 \pm 1.8 \times 10^7$ | - |
| U Fabric | $3.5 \times 10^5 \pm 3.9 \times 10^5$ | ≈ 0 % | $2.5 \times 10^6 \pm 5.1 \times 10^5$ | ≈ 0 % | $4.3 \times 10^6 \pm 1.3 \times 10^6$ | ≈ 0 % |
| HS Fabric | 0.0±0.0 | 100 % | 0.0±0.0 | 100 % | 0.0±0.0 | 100 % |

The titers obtained by contacting the phage phi 6 with the U Fabric are similar to the control as expected, within the experimental uncertainty (Table 1). However, the HS Fabric exhibited a potent phage inactivation (100% viral of viral inhibition).

Viral particles expelled by infected individuals during breathing, speaking, coughing or sneezing will come to the developed antiviral fabric in droplets containing high amounts of water.^48^ Therefore, since hand soap is very soluble in water, we believe that viral inactivation will occur very fast as confirmed by the antiviral results shown in Figure 4 and Table 1.

#### 2.2.2. Tests with SARS-CoV-2

The median tissue culture infectious dose (TCID50/mL) test of SARS-CoV-2 showed a significant reduction of infectious titers after being in contact with the HS Fabric for 1 minute from 5,16 or 5,21 to 3,46 (see Figure 5).

**Figure 5.**
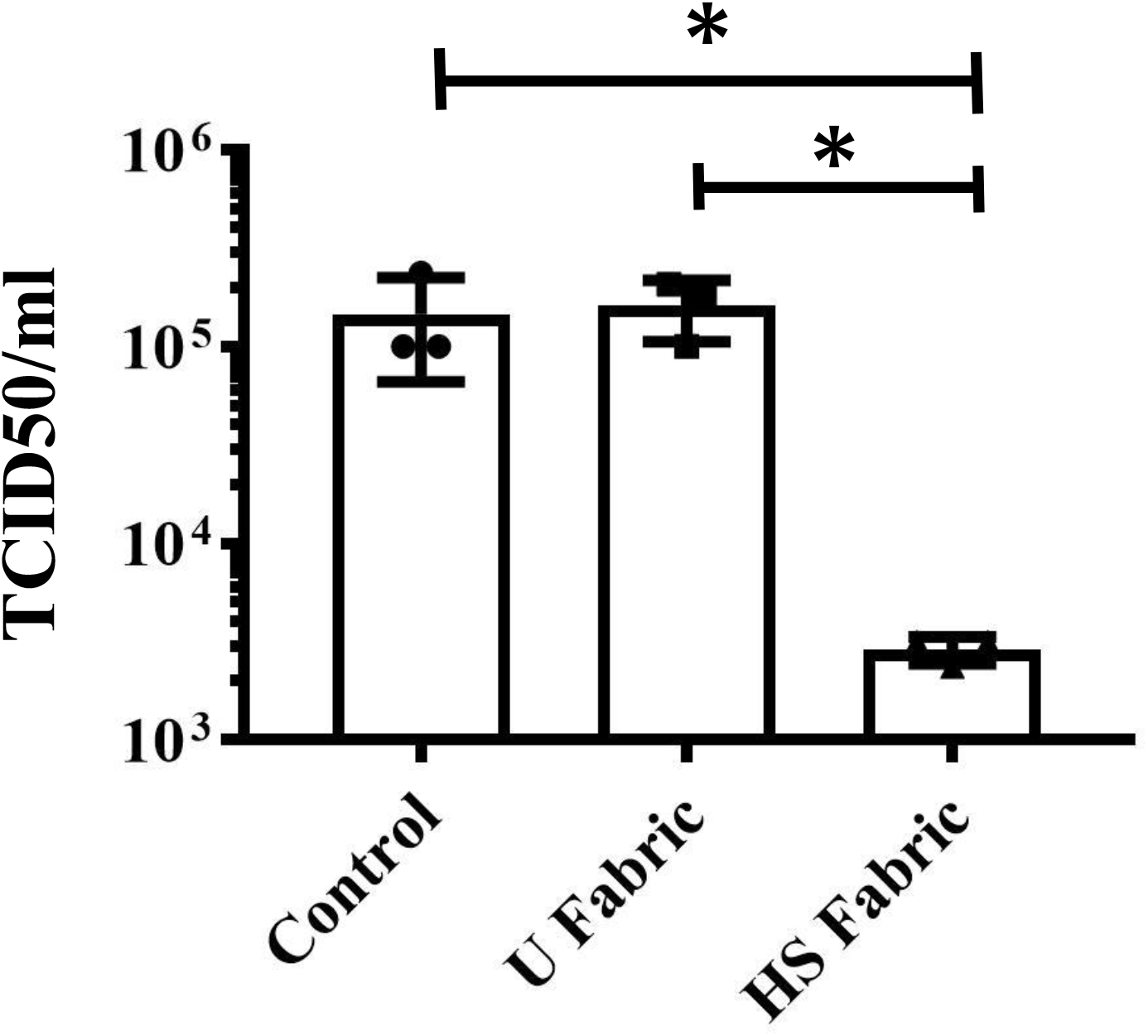
Infectious titers of SARS-CoV-2 after 1 minute of contact with the untreated fabric (U Fabric) and the fabric functionalized with solidified commercial hand soap (HS Fabric) and control (without being in contact with any fabric) by the TCID50/mL method. * *p* > 0.05; ns, not significant.

However, the U Fabric and control showed no reduction of titers as expected. These results clearly demonstrate the potent antiviral activity of the HS Fabric against SARS-CoV-2 even after 1 minute of contact and are in good agreement with those obtained with the enveloped bacteriophage phi 6 (see Figures 4 and Table 1). The infection titers obtained by the TCID50/mL method for SARS-CoV-2 are shown in Table 2.

**Table 2.** Infection titers and log values obtained by the double layer method for the antiviral assay performed with phi 6 and by the TCID50/mL method for SARS-CoV-2 expressed as mean±standard deviation, and log reductions and percentages of viral inactivation with respect to control:without being in contact with any fabric (control), and after one minute of contact with the untreated fabric (U Fabric), and with the fabric functionalized with the solidified hand soap coating (HS Fabric).

| Sample | SARS-CoV-2 at 1 min<br>(TCID50/mL) | SARS-CoV-2 at 1 min<br>Log (TCID50/mL) | Log<br>reduction | Inactivation<br>at 1 min |
| --- | --- | --- | --- | --- |
| Control | $1.46 \times 10^5 \pm 7.90 \times 10^4$ | $5.16 \pm 0.21$ | - | - |
| U Fabric | $1.62 \times 10^5 \pm 5.57 \times 10^4$ | $5.19 \pm 0.16$ | $\approx 0$ | $\approx 0 \%$ |
| HS Fabric | $2.90 \times 10^3 \pm 4.56 \times 10^2$ | $3.46 \pm 0.07$ | 1.7 | 98.0 % |

In this study, two different antiviral test methods have been performed considering the characteristics of each type of virus (phi 6 or SARS-CoV-2). However, these results confirm that the phage phi6 is a good biosafe viral model of SARS-CoV-2 against hand soap other than quaternary ammonium salts such as benzalkonium chloride.^49^ Thus, this suggests that this viral model could be very useful for researchers working in this field and not having access to a biosafety level 3 laboratory to test other screening agents with potential antiviral activity against SARS-CoV-2 or other enveloped RNA viruses such as influenza.^50,51^

### 2.3. Cytotoxicity tests of the antiviral face mask

The developed antiviral non-woven fabric was used to fabricate an antiviral face mask (see Figure 6).

**Figure 6.**
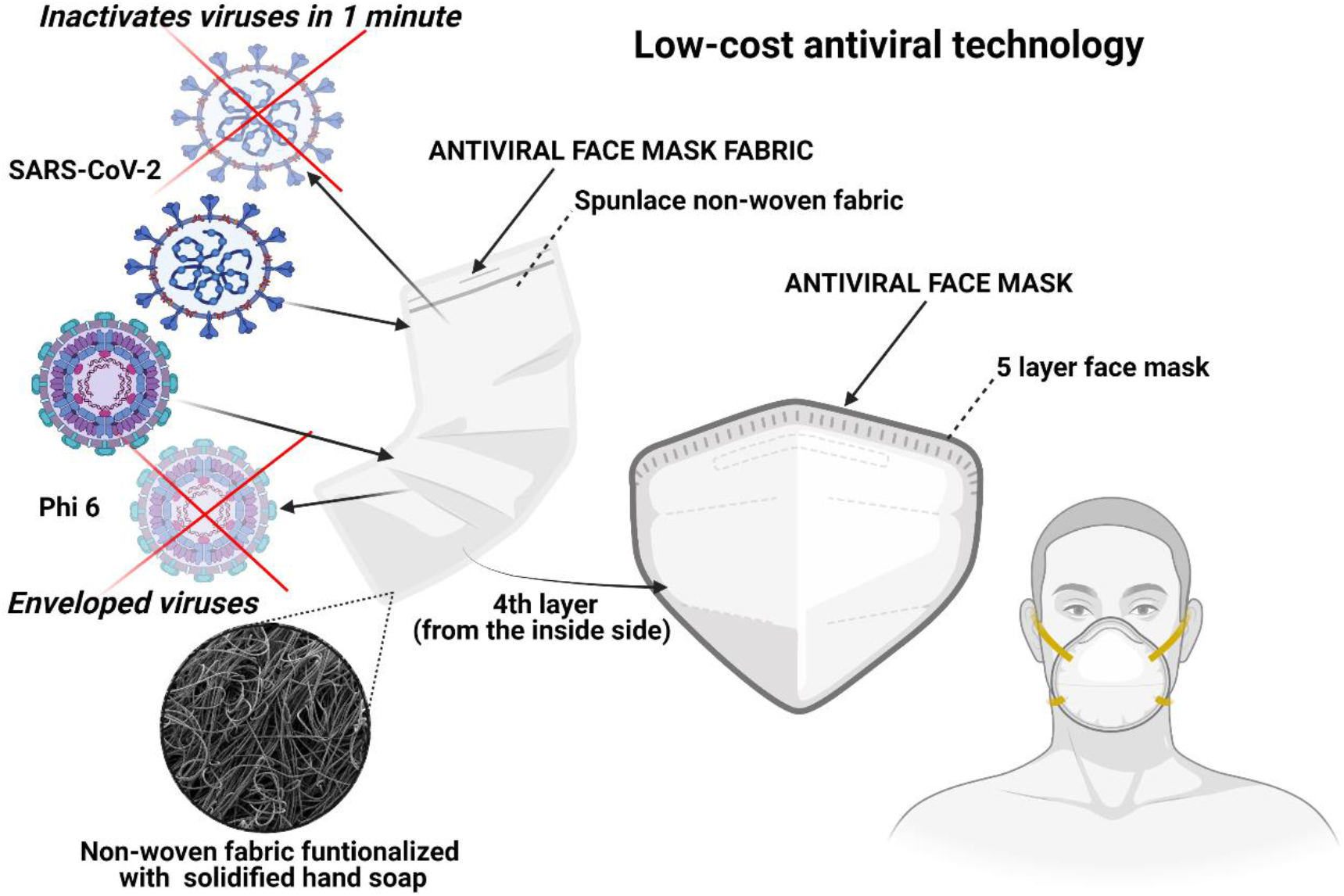
Low-cost face mask produced with an antiviral non-woven fabric functionalized with solidified hand soap capable of inactivating enveloped viruses such as SARS-CoV-2 and phi 6 in one minute of contact.

In order to avoid any direct contact of the antiviral fabric with the face of the user, the fabric was placed in the 4^th^ layer from the inside side in a commercial WottoCare (Quanzhou Huanda Bags CO. LTD, Quanzhou, China) face mask of 5 layers. A similar face mask without placing the antiviral fabric inside was used as control material. Thus, cytotoxicity tests were performed with these face mask produced with (FM) and without the antiviral non-woven fabric (FMC). Figure 7 shows that the antiviral face mask produced in this study does not produce any cytotoxic effect in human keratinocyte cells.

**Figure 7.**
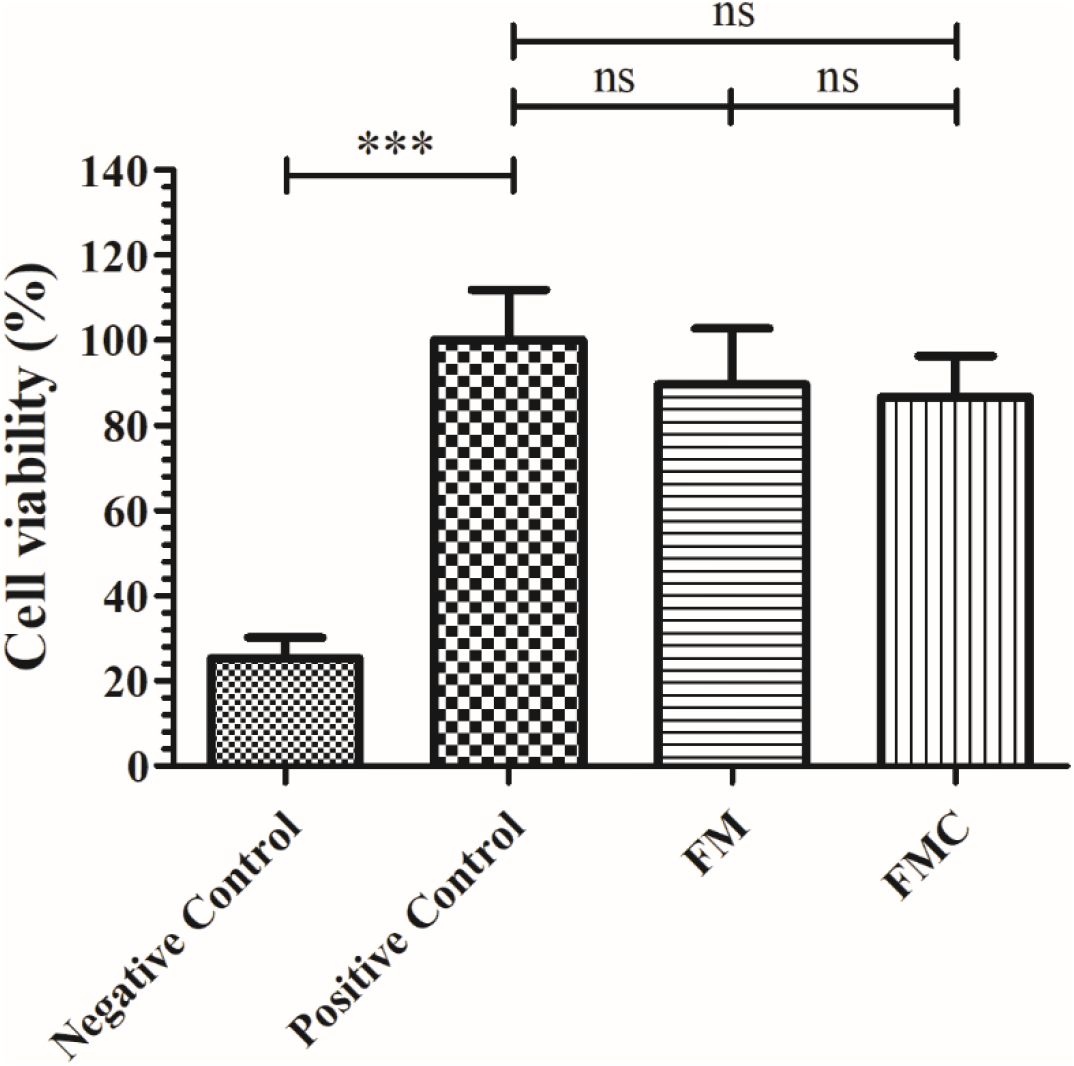
The 3-[4, 5-dimethylthiazol-2-yl]-2, 5 diphenyl tetrazolium bromide (MTT) cytotoxicity assay of extracts obtained from the 5-layer face mask with (FM) and without (FMC) the antiviral non-woven fabric functionalized with solidified hand soap, positive and negative controls cultured in the presence of human keratinocyte HaCaT cells at 37 °C. *** *p* > 0.001; ns, not significant.

The antiviral action of hand soap is mainly attributed to the surfactant molecules present in the hand soap and its mechanisms of action are attributed to membrane rupture, simple elution mechanism or viral entrapment.^52^ Thus, in this study, we have developed a new low-cost and non-cytotoxic face mask with antiviral properties to reduce COVID-19 infections in both senses, from and to the individuals wearing the mask with the HS fabric. The non-woven fabric has been developed here by a simple, low-cost, reliable and reproducible dip-coating method that uses commercial nonwoven fabric with cylindrical polymer filaments functionalized with solidified hand soap physically absorbed onto the surface of each filament.^53^ The same fabrication procedure could be applied to any type of face mask and other infection prevention clothing such as caps, scrubs, shirts, trousers, disposable gowns, overalls, hoods, aprons and shoe covers, which can be biodegradable or not to inactivate enveloped viruses such as SARS-CoV-2. This easy fabrication procedure opens an increasing number of possible applications in need of new antimicrobial approaches. Nonetheless, despite the results achieved, further research is needed to ensure that this technology is completely safe for the production and commercialization of this antiviral non-woven single-use fabric at a large scale depending on the specific application.

## 3. CONCLUSIONS

An antiviral fabric has been developed with a non-woven fabric containing polymer filaments that were functionalized with solidified commercial hand soap. This antiviral non-woven fabric was capable of inactivating enveloped viruses such as SARS-CoV-2 and phi 6 in just one minute of contact. These results demonstrate that the phage phi 6 can be successfully used as viral surrogate of SARS-CoV-2. This antiviral fabric was produced by a reproducible, fast, and economic procedure. Furthermore, we have shown how this antiviral fabric can be used for the fabrication of antiviral face masks with no toxic effects in human keratinocyte cells. The applications of this antiviral non-woven fabric are immense in the infection prevention clothing industry. Therefore, this technology could reduce significantly the COVID-19 global spread and help in future pandemics because it can be fabricated at a very low cost, especially in low and middle-income countries.

## 4. EXPERIMENTAL SECTION

### 4.1. Dip-coating with a commercial hand soap

Commercial non-woven spunlace face mask fabric from NV EVOLUTIA were cut in the form of discs (*n*=*6*) of approximately 10 mm in diameter to be treated by the dip-coating method.^45^ These fabric discs were immersed in a very diluted (1 % *w/v*) aqueous solution of a commercial liquid hand soap (KYREY dermo, Laboratorios Forenqui, S.A., Picassent, Valencia, Spain) for 30 minutes at 23±1°C to achieve gravimetrically a very low content of dry solidified hand soap 0.57±0.03% *w/w* (HS Fabric). The KYREY dermo healthcare product has been dermatologically tested and presents a common chemical composition approved by the European authorities. Thus, according to the manufacturer, this commercial liquid hand soap is composed of many ingredients: aqua, sodium laureth sulfate, sodium chloride, cocamidopropyl betaine, glycerin, cocamide dea, disodium edta, propylene glycol, styrene/acrylates copolymer, lactic acid, parfum, benzyl alcohol, limonene, linalool, DMDM hydantoin, imidazolidinyl urea, methylchloroisothiazolinone, methylisothiazolinone and sodium benzoate. Some of these compounds have shown antiviral properties against SARS-CoV-2 such as the surfactant sodium laureth sulfate ^54^ and sodium chloride.^55^ Other compounds such as cocamidopropyl betaine or sodium benzoate have been reported as active or inactive ingredients of commercial healthcare products, respectively.^44^

Time and concentration were adjusted to obtain a dip-coating treatment with the optimal antiviral properties and the lowest amount of hand soap. The functionalized fabrics were subsequently dried at 60°C for 48 hours to constant weight to solidify the physically absorbed hand soap onto the surface of the filaments of the non-woven fabrics. Discs (*n*=*6*) prepared from the untreated non-woven fabric (U Fabric) were prepared as reference material. Sterilization was subsequently performed by ultraviolet radiation one hour per side.

### 4.2. Fabric morphology and characterization

A Zeiss Ultra 55 field emission scanning electron microscope (FESEM, Zeiss Ultra 55 Model) was used with an accelerating voltage of 3 kV. Porous morphology of the treated and untreated non-woven fabric was observed at a magnification of 50 times or 1000 times. The fabric samples were prepared to be conductive by platinum coating with a sputter coating unit. The differential pressures of the U Fabric and HS Fabric were measured according to the standard EN14683:2019+AC:2019, as a measure of breathability. Thus, five specimens of 4.9 cm^2^ were used in these tests performed at 22±1°C, 30% relative humidity and with 8±0.2 L/min air flow.

### 4.3. Antiviral test using enveloped phage phi 6

The Gram-negative *Pseudomonas syringae* (DSM 21482) from the Leibniz Institute DSMZ-German collection of microorganisms and cell cultures GmbH (Braunschweig, Germany) was grown in solid Tryptic Soy Agar (TSA, Liofilchem), and after that in Liquid Tryptic Soy Broth (TSB, Liofilchem) at 25 °C and a speed of 120 r.p.m. The enveloped phage phi 6 (DSM 21518) was propagated according to the specifications provided by the Leibniz Institute DSMZ-German collection of microorganisms and cell cultures GmbH (Braunschweig, Germany).

Double-stranded RNA extraction and quantification from the phi 6 virus were performed to test whether viral particles remain attached to the U Fabric and HS Fabric materials or not compared to control before the antiviral assays to avoid false results. The amount of virus dispersed on the material discs was 50 μL of the 1/1000 dilution and left to incubate at 25°C for 5 minutes in the U Fabric and HS Fabric. The same volume of 50 μL of the 1/1000 dilution was left to incubate at 25°C for 5 minutes without being in contact with any fabric (control). After the incubation time, they were introduced into 10 mL of TSB and sonicated for 5 minutes and vortexed for 45 seconds. Thus, the next step consisted of extracting the viral RNA using the RNA extraction protocol provided by Norgen Biotek Corp (Ontario, Canada).^56^ This protocol consists of a first step of viral particle lysing to ensure that the mixture becomes transparent before proceeding to the next step. The second step consists of the viral RNA purification which concerns a series of steps: binding of this molecule to the purification column, washing of the purification column and elution of the RNA for storage avoiding degradation in a freezer at −70°C. After the RNA extraction, the amount of RNA present in the samples was quantified using a nanodrop (ThermoScientific, Waltham, USA) and the result were expressed in ng/μL. These measurements were performed in triplicate to ensure reproducible results.

The antiviral test consisted of adding a volume of 50 μL of a phage suspension in TSB to each fabric disc at a titer of approximately 1×10^6^ PFU/mL to be incubated for 1, 10 and 30 minutes. Each fabric disc was located in a falcon tube with 10 mL TSB that was sonicated for 5 minutes at 24 °C, and subsequently vortexed for 1 minute. Serial dilutions were made with each falcon for phage titration. A volume of 100 μL of each phage dilution was mixed with 100 μL of the host strain at OD_600 nm_ = 0.5. Thus, the infective activity of the phage was determined based on the double-layer method.^57^ 4 mL of top agar (TSB + 0.75% bacteriological agar, Scharlau) with 5 mM CaCl_2_ was mixed with the phage-bacteria suspension to be finally poured on TSA plates. Then, the plates were incubated in a refrigerated oven at 25°C for 24-48 hours. The phage titer of each type of sample was calculated and expressed in PFU/mL to be compared with the control sample, which consisted of 50 μL of phage mixed with the bacteria without being in contact with any fabric and without performing sonication/vortexing. The antiviral activity was calculated in log reductions of titers at 1, 10 and 30 minutes of contact with the virus model. It was checked that the residual amounts of hand soap in the titrated samples did not interfere with the titration procedure and the sonication/vortexing treatment did not affect the infectious activity of the phage. The antiviral tests were performed in triplicate during two different days (*n*=*6*) to ensure reproducible results.

### 4.4. Antiviral tests using SARS-CoV-2

The SARS-CoV-2 (SARS-CoV-2/Hu/DP/Kng/19-027) kindly gifted from Dr. Tomohiko Takasaki and Dr. Jun-Ichi Sakuragi (Kanagawa Prefectural Institute of Public Health). The SARS-CoV-2 was stored at −80°C. A volume of 50 μL of the PBS containing SARS-CoV-2 (1.3×10^5^ TCID50/fabric) was added to each fabric, and then incubated for 1 minute at 25°C. Each fabric was vortexed in 1 mL PBS for 5 minutes at room temperature. Viral titers were measured by the TCID50 assays. TMPRSS2/Vero cells^58^ (JCRB1818, JCRB Cell Bank), which were cultured with the Minimum Essential Media (MEM, Sigma-Aldrich) supplemented with 5% fetal bovine serum (FBS) and 1% penicillin/streptomycin (P/S), were seeded into 96-well cell culture plates (Thermo Fisher Scientific). Samples were serially diluted 10-fold from 10^−1^ to 10^−8^ in the MEM containing 5% FBS and 1% P/S. Dilutions were placed onto the TMPRSS2/Vero cells in triplicate and incubated at 37°C for 96 hours. Cytopathic effect was evaluated under a microscope. TCID50 values were calculated using the Reed-Muench method. All SARS-CoV-2 infection experiments were performed at a Biosafety Level 3 laboratory (Kyoto University)

### 4.5. Toxicological study of the face mask

In order to ensure the safety and correct use of this low-cost technology, the toxicity of the face masks containing the developed antiviral non-woven fabric was studied by the Norm ISO-10993 standard recommendations for face masks. Thus, face mask discs with and without the antiviral non-woven fabric functionalized with solidified hand soap of about 1 cm in diameter were cut with a cylindrical punch and sterilized under ultraviolet light for 1 hour per side. Every disc (*n*=6) was placed into a well of a 6-well plate with 1 mL of Dulbecco’s Modified Eagle’s Medium (DMEM, Biowest SAS, France) without fetal bovine serum (FBS, Biowest SAS, France). Thus, a volume ratio of 0.1 g/mL was selected according to the ISO-10993 that recommends this rate for irregular porous materials of low density such as textiles. After incubating the discs in humidified 5% CO_2_/95% air ambient for 72 hours at 37 °C, the extracts were collected, and utilized immediately for the toxicological assay. A cell line of non-tumorigenic immortalized human keratinocyte HaCaT cells was provided from the Medical Research Institute Hospital La Fe, Valencia, Spain. Cell incubation was performed in a mixture of DMEM with 10% FBS, 100 units/mL penicillin (Lonza, Belgium) and 100 mg/mL streptomycin (HyClone, GE Healthcare Life Sciences), at 37 °C and 5% CO_2_. The effect of the treatment with the face mask extracts on cell viability was measured by the 3-[4, 5-dimethylthiazol-2-yl]-2, 5 diphenyl tetrazolium bromide (MTT) assay. The HaCaT cells were planted at 10^4^ cells/well onto 96-well plate. After incubation for 24 hours at 37 °C, the medium in each well was replaced with 100 μL of face mask disc extracts. The medium was also replaced with 100 μL of the same medium used to produce the face mask disc extracts (positive control), as well as 100 μL of 1000 μM zinc chloride (≥ 97.0%, Sigma Aldrich) solution as negative control because this concentration is highly toxic for the HaCaT cells.^59^ Cell incubation was performed with 5 mg/mL MTT in each well for 3 h. Thus, the formazan crystals were dissolved in 100 μL dimethyl sulfoxide (DMSO, Sigma Aldrich) at ambient temperature and the absorbance was read at 550 nm on a microplate reader (Varioskan, Thermo Fisher).

### 4.6. Statistical analysis

Student’s t-test was performed for pair comparisons and one-way analysis of variance (ANOVA) for multiple value comparisons followed by Tukey’s posthoc test (\**p* > 0.05, \*\*\**p* > 0.001) on GraphPad Prism 6 software.

## Author contributions

Á.S-A conceived the idea of this work, wrote the original draft manuscript and performed the figure design. A.C-V, and A.T-M. contributed equally to this work. Methodology, validation and formal analysis: M.M, K.T and Á.S-A; software: K.T and Á.S-A; investigation: M.M, A.T-M, A.C-V., Y.M, T.N, K.T and Á.S-A; resources: M.M, K.T and Á.S-A; data curation: A.T-M, A.C-V., K.T and Á.S-A; writing-review and editing: A.T-M, A.C-V., M.M, K.T and Á.S-A; supervision: M.M, K.T and Á.S-A; project administration: K.T, T.N and Á.S-A; funding acquisition: K.T and Á.S-A.

## Notes

The authors declare no competing financial interest.

## ACKNOWLEDGMENTS

The authors would like to express their gratitude to the Fundación Universidad Católica de Valencia San Vicente Mártir for their financial support through the Grant 2020-231-006UCV and to the Ministerio de Ciencia e Innovación (PID2020-119333RB-I00 / AEI / 10.13039/501100011033) (awarded to Á.S-A). This research was also supported by grants from the Japan Agency for Medical Research and Development (AMED) (20fk0108270h0001, 20fk0108263s0201). This work was supported by Joint Usage/Research Center program of Institute for Frontier Life and Medical Sciences Kyoto University. We would like to thank Dr. Yoshio Koyanagi and Dr. Kazuya Shimura (Kyoto University) for setup and operation of the BSL-3 laboratory.

## ABBREVIATIONS

COVID-19: coronavirus disease 2019
SARS-CoV-2: Severe Acute Respiratory Syndrome Coronavirus 2
U Fabric: untreated fabric
HS Fabric: fabric functionalized with solidified commercial hand soap
FM: face mask with antiviral fabric
FMC: face mask with antiviral fabric
TSA: Tryptic Soy Agar
TSB: Liquid Tryptic Soy Broth
PFU/mL: plaque-forming units per mL
TCID50: median tissue culture infectious dose
FBS: fetal bovine serum
MEM: Minimum Essential Media
DMEM: Dulbecco’s Modified Eagle’s Medium
MTT: 3-[4, 5-dimethylthiazol-2-yl]-2, 5 diphenyl tetrazolium bromide
DMSO: dimethyl sulfoxide

